# Unraveling the inverse latitudinal gradient: Environmental and geographic influences on plant diversity in South American coastal lomas

**DOI:** 10.1101/2024.10.23.619859

**Authors:** Fiorella N. Gonzales, Holger Kreft, Luis Arapa, Dylan Craven

## Abstract

Water availability is widely recognized for its importance in shaping plant diversity gradients globally. However, even in arid ecosystems such as drylands, a remarkable diversity of plant species can persist. Yet, the environmental, topographic, and geographic drivers that have structured plant diversity gradients within drylands over evolutionary time, and their relative importance, are poorly understood, particularly for understudied ecosystems such as coastal lomas. Here, we evaluated the geographic distribution and drivers of taxonomic and phylogenetic diversity of plants across the coastal lomas of South America, which are isolated fog-fed vegetation oases within the Peruvian and Chilean coastal desert. We collated the most comprehensive data set to date of 72 coastal lomas, and used generalised linear models to assess latitudinal diversity gradients, and evaluated the relative importance of environmental, topographic, and geographic drivers of plant taxonomic and phylogenetic diversity. Notably, we found that both plant species richness and standardized mean nearest taxon distance (MNTD) decreased linearly towards the Equator, while standardized phylogenetic diversity (PD) and mean phylogenetic distance (MPD) exhibited non-linear relationships with latitude, peaking in the hyper-arid core of the Atacama Desert. Our results suggest that current climate, environmental heterogeneity, and geographical factors are the primary drivers of plant diversity. Plant species richness increased with cloud cover, slope, and area, and decreased with soil pH, while standardized PD increased with aridity, elevation, and the human footprint and decreased with distance to the coast. Standardized MNTD increased with elevation, and decreased with increasing area and temperature; standardized MPD increased with slope, and decreased with increasing aridity and distance to the coast. Our results suggest that environmental filtering is not the only macroecological process acting on plant diversity across coastal lomas, and has often been counterbalanced at different points in evolutionary time by factors associated with environmental heterogeneity, possibly reflecting the presence of climate refugia and diverse habitats that promote species coexistence. Also, our results offer empirical evidence supporting an inverse latitudinal gradient in plant diversity across coastal lomas, which emerges as a consequence of multiple factors that influence water availability, both directly and indirectly. Over evolutionary time, these factors may have significantly contributed to shaping the current structure and composition of coastal lomas.

## 1. Introduction

Despite their aridity, drylands are home to 20% of global plant diversity (White & Nackoney, 2003) and provide critical ecosystem services to more than 38% of the world’s population (Reynolds et al., 2007; United Nations Development Program UNDP, 2014). Yet, the biodiversity of these ecosystems is highly vulnerable to global change drivers, principally desertification and land-use change (Maestre et al., 2012), which may have cascading effects across different taxonomic groups and ecosystem functioning (Maestre et al., 2016). Understanding the macroecological and macroevolutionary processes that generate - and maintain - biodiversity in the most arid ecosystems globally may therefore provide insights into the potential impacts of aridification of other ecosystems (Tariq et al., 2024).

The long history of drylands is coupled with the history of its major plant lineages, with major dryland clades having diversified more or less in synchrony between the Late Miocene (11.63–5.33 Ma) and the Early Pliocene (5.3–3.6 Ma) (Maestre et al., 2021).

Drylands show a heightened recent diversification -showing short branch length- resulting in a lower number of plants compare to extratropical moist biomes (Neves et al., 2021). An example of this is Caatinga dry forest, which is an important center of diversification of Cactaceae family (Ortega-Baes & Godínez-Alvarez, 2006). Unlike other drylands (e.g., Luebert, 2021; Maestre et al., 2021; Saiz et al., 2018; Ulrich et al., 2016; Wang et al., 2017) biodiversity patterns of coastal lomas – isolated vegetation oases distributed within the Peruvian Chilean coastal desert – and its macroecological and macroevolutionary drivers are poorly understood (Gonzales et al., 2023).

Coastal lomas are sustained by advective fog deposition that occurs on steep coastal slopes (< 1000 m a.s.l) are distributed between 200 and 1200 meters in elevation, within the hyper-arid desert. In addition, coastal lomas are influenced by El Niño Southern Oscillation (ENSO) events, which trigger periods of rainfall (Dillon & Rundel, 1990, Cereceda et al., 2002). Furthermore, ENSO events impact the occurrence of marine fog across the coastal desert. During strong to extreme ENSO events, marine fog becomes more frequent during the summer months at ∼20°S (del Río et al., 2018), which have also been associated with the intrusion of fog with more precipitable water (Eichler & Lodoño, 2013). Coastal lomas vegetation is dominated by a mixture of annual and perennial grasses, herbs, shrubs, and small populations of trees (Dillon et al., 2003). Tree species like *Myrcianthes ferreyrae* are endemic to the coastal desert and occur in just a few lomas (Gonzales & Villasante, 2019). Fog condensation increases with altitude until the upper limit reaches the regional temperature inversion (Schemenauer & Cereceda, 1993), reaching its maximum at intermediate elevations (Rundel & Mahu, 1976), but can vary considerably, depending on proximity to the ocean, elevation, local topographic relief, and aspect (Larrain et al., 2002). The extreme climate of the coastal desert also is thought to have partitioned the flora of the coastal lomas into three distinct assemblages (Dillon et al., 2003; Gonzales et al., 2023), with the hyper-arid zone between between 18° and 22 S° activing as a dispersal barrier (Dillon, 2005; Rundel et al., 1991).

Several ecological mechanisms have been hypothesized to elucidate diversity patterns in drylands (see Li et al., 2013; Bergholz et al., 2017; Maestre et al., 2021), the principal among which being aridity (i.e., the ratio of precipitation (P) to potential evapotranspiration (PET)) (e.g., Berdugo et al., 2020; Liu et al., 2019; Maestre et al., 2016; Wion et al., 2020). Generally, plant species richness in drylands is strongly influenced by water availability (Li et al., 2013), sometimes even increasing with a decrease in water availability (Wang et al., 2018). In coastal lomas, water supply comes from two main sources: (i) ENSO events, whose brief periods of heavy rains stimulate plant growth and increase recruitment and diversity (Dillon & Rundel, 1990; Jiménez et al., 1999; Muñoz-Schick et al., 2001; Pinto et al., 2001; Rundel et al., 2007; Tovar et al., 2019) resulting in marked differences in species composition across the distribution of coastal lomas (Dillon, 1997; Manrique et al., 2014; Pino & Luebert, 2009), and (ii) coastal fog, which creates vegetation bands that are associated with higher levels of humidity that vary in elevation (Larrain et al., 2002; Schemenauer & Cereceda, 1993). At local spatial scales, plant species richness and net primary productivity NPP increase with elevation, which is usually associated with higher levels of humidity in coastal lomas (Ávila & Guevara, 2006; Cuya, 2016; Muenchow et al., 2013; Sotomayor & Jiménez, 2008). Distance to the coast is another proxy for water availability that may positively affect plant diversity (Moat et al., 2021), as proximity to the coast implies higher water availability, with steeper coastal hills intercepting more fog than flatter areas where the stratus layer of fog dissipates. Other factors, such as soil conditions (e.g., soil C:N, soil pH) and temperature (e.g., mean annual temperature), may mitigate or exacerbate the effects of water availability on plant diversity (Nunes et al., 2019; Zhang et al., 2024). However, the joint impacts of environmental drivers on plant diversity patterns across coastal lomas have yet to be assessed, as previous efforts have been limited by the data availability (Manrique et al., 2014).

In parallel with mean environmental conditions (see above), their variability can also influence biodiversity patterns across coastal lomas. In particular, environmental heterogeneity is crucial for maintaining the richness of rare species in drylands (Liu et al., 2019). Environmental heterogeneity resulting from small-scale topographic variability creates microclimatic conditions that buffer communities from the direct effects of aridity (e.g., Keppel et al., 2018) and prevents the loss of less stress-tolerant individuals or species (Stotz et al., 2021). Moreover, the climate that plants directly experience can be mediated by ‘nurse plants’, highlighting the role of plant cover in facilitating species coexistence - and, indirectly, ecosystem functioning (Beugnon et al., 2024) - in extreme environments such as deserts (Bertness & Calloway, 1994; Maestre et al, 2009; Sotomayor & Drezner, 2019). Lastly, small-scale microclimate variability in dryland soils contribute to the maintenance of community evenness and the type of species distribution (Ulrich et al., 2016). Because average environmental conditions fail to capture the effects of abiotic or biotic facilitation, it is therefore crucial to also consider environmental heterogeneity to deepen current understanding of the drivers of biodiversity patterns in water-limited ecosystems such as coastal lomas.

As expected, the species-area relationship hypothesis (Rosenzweig, 1995) holds in coastal lomas ecosystems (Arana, 2019), and is primarily influenced by spatial aggregation, species evenness and plant cover (DeMalach et al., 2019, McGlinn et al., 2019). Unlike in other ecosystems, latitudinal diversity gradients (LDGs) in drylands may be less pronounced due to harsh environmental factors (e.g., higher aridity, low annual precipitation). In the case of coastal lomas, plant species richness does not vary linearly with increasing latitude, possibly due to high levels of endemism and climatic barriers that maintain diversity at similar levels across the distribution of coastal lomas (Manrique et al., 2014). However, the proximate drivers underlying the spatial variation in plant diversity across coastal lomas –and their relative importance – have yet to be explored within the frameworks of other relevant macroecological theories, particularly that of water-energy dynamics (Hawkins et al., 2003; Vetaas, 2019).

In coastal lomas, taxonomic diversity has been the most frequently used measure of diversity, particularly at local spatial scales (Gonzales et al., 2023). Yet, other components of biodiversity that can provide complementary insights to ecological mechanisms shaping community assembly, such as phylogenetic diversity and structure metrics such as Faith’s phylogenetic diversity (Faith’s PD), which is the sum of the branch lengths of the phylogenetic tree (Faith, 1992), mean pairwise distance between species (MPD) and the pairwise distance between the closest relatives in an assemblage (MNTD) (Webb et al., 2002). As Faith’s PD is usually closely and positively related with richness (Cadotte & Davies, 2016), the use of null models that maintain species richness constant while randomizing phylogenetic relationships facilitates comparison across assemblages (Kembel & Hubbell, 2006; Mazel et al., 2016). Consequently, negative values of standardized phylogenetic diversity and structure metrics reflect phylogenetic clustering (i.e., when species are more closely related than expected by chance), while positive values of these same metrics reflect phylogenetic overdispersion (i.e, when species are more distantly related than expected by chance; Mazel et al., 2016). Evaluating phylogenetic metrics in coastal lomas may yield new insights to underlying macroevolutionary processes, by identifying the relative importance of environmental filtering, limiting similarity and niche conservatism, and how they may shift over evolutionary time (Hardy & Senterre, 2007; Mazel et al., 2016; Tucker et al., 2016).

Phylogenetic clustering occurs where evolutionary distances between species are lower than expected by chance, and can result from environmental filtering and indicates that environmental conditions act as a filter selecting for species with certain trait values, or that a particular region has rapid speciation rates or slow extinction rates (Kraft et al., 2015; Wiens & Graham, 2005). In contrast, limiting similarity (or phylogenetic overdispersion) occurs where evolutionary distances between species are greater than expected by chance, and indicates that niche partitioning and biotic processes decrease the evolutionary similarity of co-occurring species (Burns & Strauss, 2011; Webb et al., 2002). Because of the fundamental role played by aridity in shaping biodiversity patterns in other drylands (e.g., Maestre et al., 2021), we expect that environmental filtering has been the dominant process in determining plant diversity patterns across coastal lomas over evolutionary time, largely due to the rapid diversification in some plant linneages since the Late Pliocene (Maestre et al., 2021; Neves et al., 2021).

Here, we examine the environmental, topographic, and geographic drivers of plant diversity and its underlying ecological mechanisms across coastal lomas in South America. Our main goal was to identify the environmental, topographic and geographical factors that determine plant taxonomic and phylogenetic diversity across coastal lomas. Because standarized Faith’s PD and taxonomic diversity are usually correlated, we expected that both metrics will exhibit an inverse latitudinal gradient, as coastal lomas with more energy (i.e., annual potential evapotranspiration (PET), actual evapotranspiration (AET) can support more diverse assemblages (Currie, 1991; Cerezer et al., 2022), but standardized MNTD, a terminal metric that captures phylogenetic structure at shallower evolutionary timescales, will exhibit a non-monotonic latitudinal gradient due to recent mid-latitude plant diversification along the coastal desert, and standardized MPD, a basal metric associated with phylogenetic structure at deeper evolutionary timescales, will exhibit a non-linear relationship with latitude due to increasing aridity between 18°and 22°S, and increasing humidity towards the northern distribution of coastal lomas (Rundel et al., 1991). We further expected that environmental filtering, likely associated with water availability, will constrain plant diversity (taxonomic and Faith’s PD) in more arid areas across the distribution of coastal lomas. We therefore hypothesized that taxonomic and Faith’s PD will increase with area, environmental conditions, i.e., mean annual temperature (MAT), mean annual precipitation (MAP), cloud cover, soil properties, and environmental heterogeneity (slope and elevation), and decrease with aridity and distance to the coast (Arana, 2019; Moat et al., 2021). In coastal lomas further from coast with higher MAT and aridity yet lower MAP, cloud cover, lower soil N and pH and environmental heterogeneity, we expected that standardized MNTD and MPD will exhibit phylogenetic clustering (i.e., negative standardized MNTD and MPD values, Swenson & Enquist, 2009), while those closer to the coast with lower MAT and aridity, and higher MAP, cloud cover, soil N and pH and environmental heterogeneity (slope and elevation) will exhibit phylogenetic overdispersion or divergence (i.e., positive standardized MPD and MNTD values; Swenson & Enquist, 2009). Finally we hypothesized that as elevation increases in coastal lomas, aridity tends to decrease due to changes in advective fog flux (see Cereceda et al., 2002, Rundel et al., 1991). That is, we expected that the phylogenetic diversity patterns across coastal lomas will be determined by different underlying ecological processes, i.e., limiting similarity or environmental filtering, largely due to shifts in aridity and its proxies over evolutionary timescales.

## 2. Methods

### 2.1 Study area

We evaluated the 72 coastal lomas, which extend through the Peruvian coastal desert and the Atacama Desert (hereinafter called coastal desert), extending continuously over 3,500 km from 7° to 30° S (Tago-Nakazawa & Dillon, 1999) (Figure 1). The climate of the coastal desert is characterized by extreme aridity, short periods of heavy rainfall, and relatively high temperatures in the northern parts of the coastal desert, bringing wet tropical conditions which are associated with ENSO events and the remarkable temperature homogeneity along the entire latitudinal extent of the coastal desert (Rundel et al., 1991). According to the climatic classification of Koppen-Geiger, the study region belongs to tropical and sub-tropical desert climate (“Bwh”; Beck et al., 2018). Coastal lomas have two distinct seasons, a dry season and a humid season (mostly between May and October), that determine changes in their productivity.

**Figure 1.**
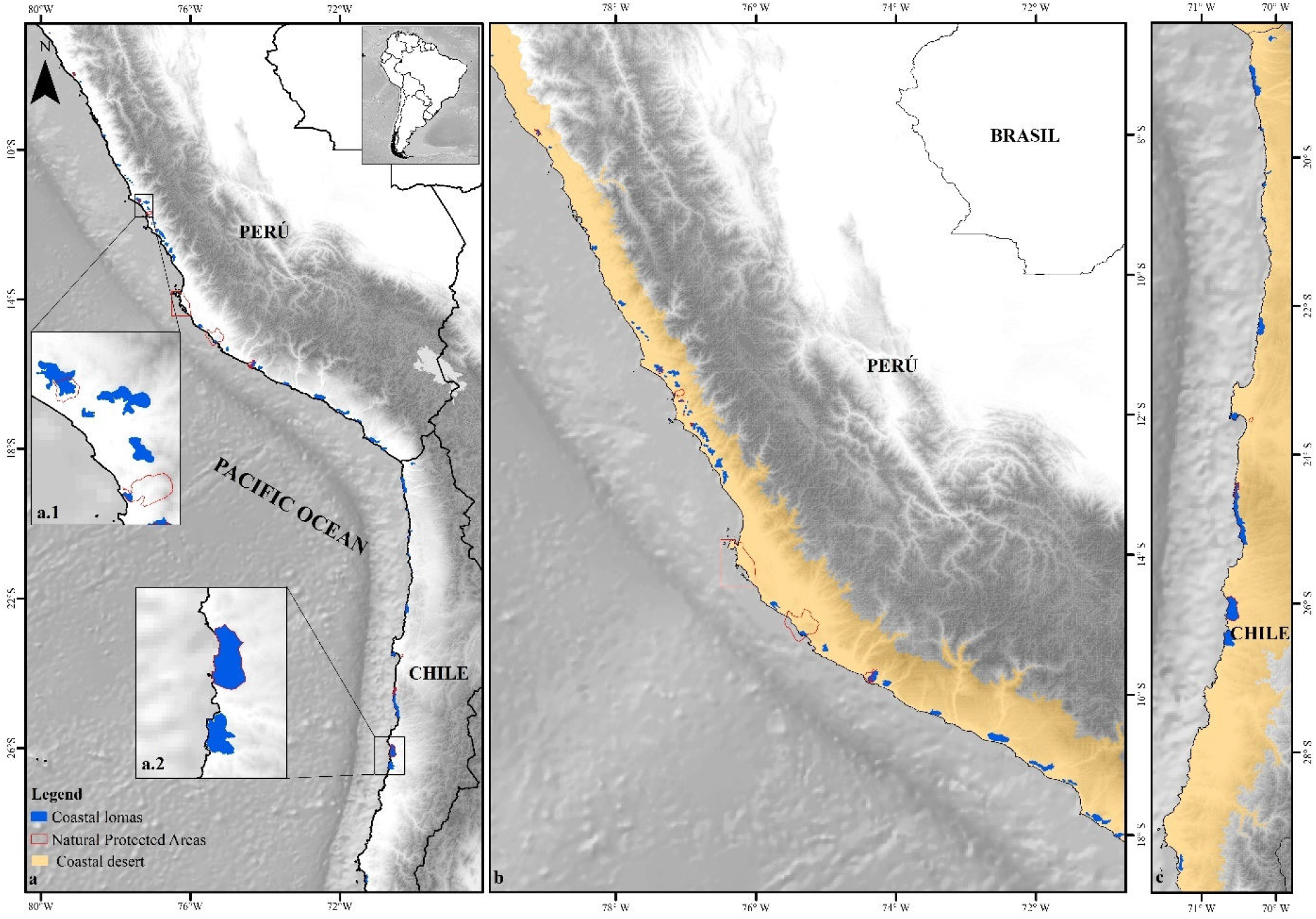
(a) Locations of coastal lomas (n = 72) across the Peruvian Chilean coastal desert of South America. Examples of the isolated distribution of coastal lomas in Lima, Peru (a.1) and in Antofagasta, Chile (a.2). In (b) and (c), the distribution of coastal lomas in Peru and Chile respectively. Coastal lomas are indicated in blue, natural protected areas are indicated with red lines, and the coastal desert is indicated by orange.

### 2.2 Vegetation data

We compiled a comprehensive database of the flora of coastal lomas that covers 72 locations. We systematically reviewed Peruvian and Chilean flora based on species records in herbaria (San Marcos Museum of Natural History UNMSM-Lima, Peru; Michael Owen Dillon Institute IMOD-Arequipa, Perú; Department of Botany of the University of Concepción CONC Concepcion, Chile). Additionally, we collected species checklists of natural reserves, scientific publications, theses, online data sets (http://sacha.org/) and personal observation, which we further complemented with species records obtained from the Global Biodiversity Information Facility (GBIF). In total, we obtained 1460 records of species that occur in coastal lomas. All data references are provided in Supporting Information (Appendix S1).

### 2.3 Data preparation

#### -Taxonomic standardization

To ensure compatibility of scientific names across our data set, we standardized species names based on the Taxonomic Name Resolution Service (Boyle et al., 2013), The Plant List (National Plan List, 2022), Tropicos (Missouri Botanical Garden, 2023), USDA (United States Department of Agriculture, 2022), and the WCVP (World Checklist of Vascular Plants, 2021) using the tnrs function of the R package ‘TNRS’ (Boyle et al., 2021). After taxonomic standardization, our data set had a total of 1,365 accepted plant species.

#### -Phylogeny

We constructed a phylogenetic tree for all plant species in our data set using the seed plant phylogeny of Smith and Brown (2018) as a backbone, using the function ‘read.newick, congeneric.merge’ of the R package ‘phylotools’ (Revell, 2024). We placed a total of 1,307 species on the phylogeny, representing 95.7% of all species.

#### -Biogeographical characteristics of coastal lomas

We spatially delimited coastal lomas by combining geospatial files from the following sources: Map of Fragile Ecosystems (Peru; SERFOR, 2019), National Map of Ecosystems (Peru; MINAM, 2019), Map of Vegetation Cover (Peru; MINAM, 2015), Map of Protected Natural Areas and Private Conservation Areas (Peru; SERNANP, 2023; Arapa, 2018), Map of Cloud Oasis Ecosystems (Chile; Moat et al., 2021), Vegetation Units (Chile; Luebert & Pliscoff, 2017) and Protected Wilderness Areas (Chile; SNASPE, 2019). For coastal lomas without geospatial information, we delimited them manually using ArcGis 10.8. To ensure the quality of geospatial files, we checked them against species records with spatial coordinates. On average, coastal lomas cover 5264.07 hectares, ranging from 77. 26 ha to 44862.44 ha.

#### -Environmental conditions

Based on relevant literature (e.g., König et al., 2017; Kreft et al., 2007; Testolin et al., 2020; Wilson & Jetz, 2016) and variables related to macroecological hypotheses that explain diversity patterns in drylands (Appendix S2), we selected an array of environmental variables. We obtained mean annual daily mean temperature (bio1), isothermality (bio3), annual precipitation (bio12), precipitation of the wettest month (bio13), precipitation of the driest month (bio14), mean monthly climate moisture index (cmi_mean), net primary productivity (NPP), and aridity, i.e. the differences between annual precipitation and potential evapotranspiration, from CHELSA 2.1 (Karger et al., 2017). As cloud cover is a proxy for fog, which is important for sustaining coastal lomas (e.g., Weberbauer, 1911; Ferreyra, 1960), we obtained annual cloud cover and cloud cover for August, which is usually the wettest month (EarthEnv, Wilson & Jetz, 2016). We derived average soil pH and N for each coastal loma, using values estimated at a depth of 5 and 15 cm from SoilGrids 1Km (Hengl et al., 2014) given the effects of soil properties on plant diversity patterns in drylands globally (Ulrich et al., 2014). We also included the human footprint index (Venter et al., 2009), because coastal lomas have been historically threatened by anthropogenic activities (Trinidad et al., 2012; Gonzales et al., 2023). All data sources had a spatial resolution of resolution 1°x1°, i.e. approximately 1 km^2^. We calculated average values for all climatic variables for each coastal loma.

#### -Topographic and geographical factors

For each coastal loma, we calculated aspect (including sine and cosine), topographic roughness, elevation, and slope following Amatulli et al. (2018) as proxies for environmental heterogeneity. As the species diversity of fragmented habitats such as coastal lomas is often associated with area and isolation (Arana, 2019), we also calculated the area (km^2^), distance to the coast (km), and geographic isolation using the surrounding mass area index following Weigelt and Kreft (2013) for each coastal loma. As for the environmental variables, we calculated topographic and geographic variables as the average for each coastal loma.

### 2.4 Data analysis

#### -Taxonomic and phylogenetic diversity

We estimated taxonomic species richness as the number of species found to occur within each coastal loma. We estimated phylogenetic diversity using three indices that target different components of evolutionary history: (i) Faith’s PD (PD; Faith, 1992), which is the sum of all branch lengths and thus represents lineage diversity of an assemblage, while (ii) mean pairwise distance (MPD; Webb et al., 2002) and (iii) mean nearest taxon distance (MNTD; Webb, 2000) emphasize different aspects of the phylogenetic structure of assemblages. MPD is the mean of the pairwise distances from each taxon to each other taxon within an assemblage and MNTD is the phylogenetic distance to the closest relative for each taxon and calculates the mean across all taxa (Webb, 2002). All PD metrics were calculated using the function pd, mpd, mntd of the R package ‘picante’ (Kembel et al., 2010).

To control for differences in species richness across coastal lomas, we used null models to compare observed and randomized species assemblages to evaluate differences in phylogenetic diversity. We, therefore, generated 1000 null communities for each coastal lomas using the ‘taxa.labels’ null model of the R package ‘picante’(Kembel et al., 2010). We then calculated standardized effect sizes for each metric of phylogenetic diversity using the functionsses.pd, ses.mpd, and ses.mntd of the R package ‘picante’ (Kembel et al., 2010). Values higher than 0 indicate overdispersion, i.e. species are less related than expected, and values lower than 0 indicate phylogenetic clustering i.e. species are more related than expected.

#### Statistical analysis

To evaluate the relationship between species richness and phylogenetic diversity with latitude, we used generalised additive models (GAM), because they capture non-linear patterns and are less sensitive to outliers (Hastie & Tibshirani, 1990). We tested the influence of climate and soil proprieties, environmental heterogeneity, and geographical factors on the variation of taxonomic and phylogenetic diversity across coastal lomas using generalised linear models (GLM). We restricted our analysis of phylogenetic diversity to standardized indices, because unstandardized phylogenetic diversity indices are strongly correlated with species richness while standardized phylogenetic diversity indices provide insights into ecological processes underlying taxonomic diversity patterns (Webb, 2002), such as environmental filtering and limiting similarity that are complementary to species richness patterns.

Except for latitude, we used a log10(x+1) transformation to reduce skewness and visually assess model assumptions, i.e. homogeneity of variance and normality of model residuals. We considered models with a ΔAIC value <2.0 compared to the minimum AIC value to be the most parsimonious models (Burnham & Anderson, 2004) using the R package “MuMin” (Barton, 2019). As we did not find significant differences between AIC values, we selected the best model using the ‘stepAIC’ function of the MASS package, which performs a stepwise procedure that iteratively adds or removes variables from the model, and selects the best model with the lowest AIC value (Venables & Ripley, 2022).

We fitted GAMs between richness and phylogenetic diversity index with latitude using the R package ‘mgcv’ (Wood, 2023) and GLMs using the “glm” function in the R packages ‘stats’ (R Core Team, 2023) the R packages ‘lm’ and ‘ggeffects’ (Lüdecke, 2018). To examine spatial autocorrelation, we calculated Moran’s I for model residuals using the R package ‘spdep’ (Bivand, 2022).

All analyses were conducted in R version 4.3.1 (R Core Team, 2023), using packages: ‘dplyr’ (Wickham et al., 2020), ‘tidyr’ (Wickham & Henry, 2019), ‘stringr’ (Wickham, 2019), ‘TNRS’(Boyle et al., 2021), ‘Taxonstand’ (Cayuela et al., 2021), ‘picante’ (Kembel et al., 2010), ‘ggeffects’ (Lüdecke, 2018), ‘ggplot2’ (Wickham, 2016), ‘cowplot’ (Wilke, 2019), ‘patchwork’ (Pedersen, 2020), ‘jtools’(Long, 2022),‘MASS’ (Venables & Ripley, 2002),‘car’(Fox & Weisberg, 2019) and ‘pgirmess’ (Giraudoux, 2021).

## 3. Results

### 3.1 Latitudinal diversity gradients

Our analysis showed that plant species richness (Pseudo R^2^=0.108, F=1.81, df=7.14, p- value <0.001) and standardized MNTD (Pseudo R^2^=0.134, F=2.33, df=5.38, p- value<0.001) decreased significantly and linearly towards the Equator (Fig.2). In contrast, standardized PD (Pseudo R^2^=0.137, F=2.44, df=5.17, p-value >0.001) and MPD (Pseudo R^2^=0.336, F=5.05, df=5.9, p-value >0.001) varied non-linearly with latitude, and MPD peaked at ∼23° S (Fig. 2). On average, we observed 84 (SE=9.37) plant species, and standarized PD of -0.43 (CI=-0.72, -0.13), a standarized MPD of -0.150 (CI=-0.44, 0.14) and standarized MNTD of -0.49 (CI=-0.76, -0.21). We found that none of the standardized phylogenetic diversity indices exhibited a statistically significant relationship with species richness (p > 0.05; Fig. S1).

**Figure 2.**
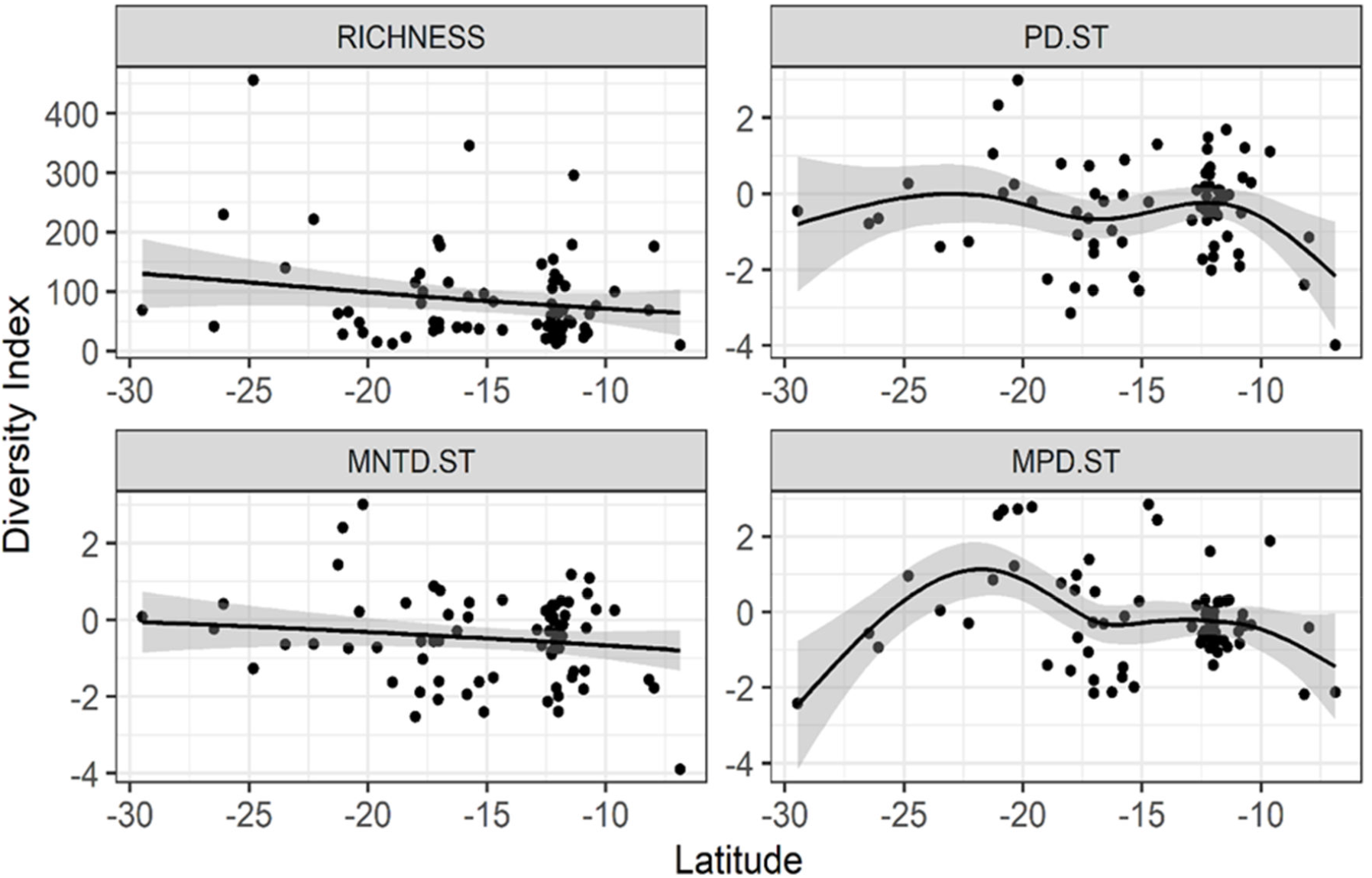
Relationships between latitude (South to North) and plant species richness or phylogenetic diversity across coastal lomas embedded within the Peruvian Chilean coastal desert (N = 72 coastal lomas). Phylogenetic diversity was calculated as MNTD, which is mean nearest taxon diversity, MPD, which is mean pairwise distance, and PD, which is Faith’s PD; these three indices were standardized with null models. Lines are fitted with generalized additive models (GAM).

### 3.2 Drivers of diversity patterns in the coastal lomas

Our results show that current climate, environmental heterogeneity, and geographical factors (area and coastal distance) jointly are the primary drivers of plant taxonomic and phylogenetic diversity. We observed that plant species richness increased in response to area, cloud cover, and slope and decreased in response to soil pH (Figs. 3 and 4, Table S1). We found that standardized Faith’s PD increased with aridity, elevation, and human footprint, yet decreased with distance to the coast (Figs. 3 and 4, Table S1). Standardized MNTD decreased with increasing area and MAT, and increased with elevation (Figs. 3 and 4, Table S1); standardized MPD increased with slope and decreased with increasing aridity and distance to the coast (Figs.3, and 4, Table S1). Our models explained a greater amount of variation for phylogenetic diversity (Pseudo R^2^ = 0.20 – 0.28) than for taxonomic diversity (Pseudo R^2^ = 0.06; Table S1).

**Figure 3.**
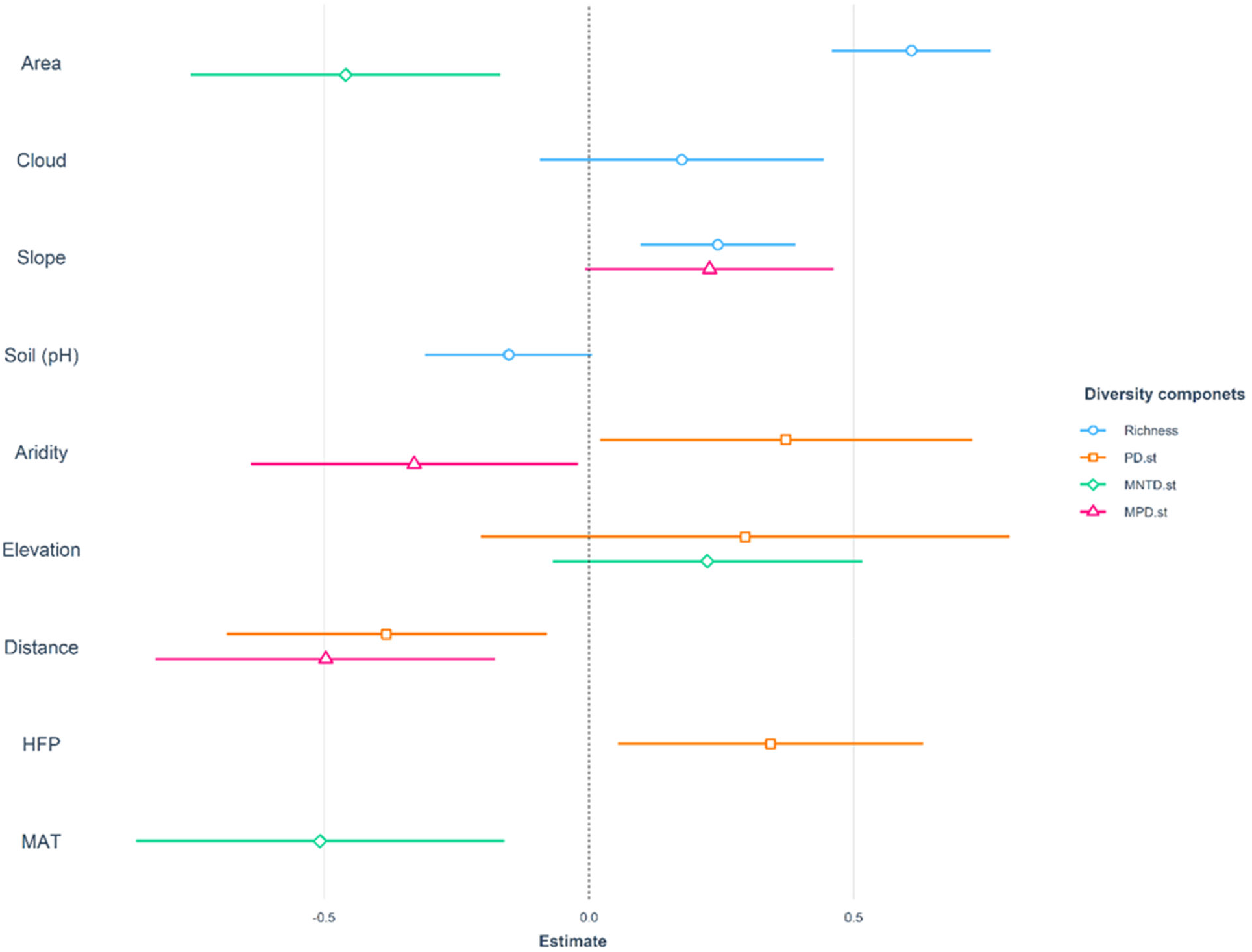
Effect sizes of abiotic and biotic predictors on plant diversity (taxonomic and phylogenetic) across coastal lomas in Perú and Chile. Richness is species richness, PD.st is standardized Faith’s phylogenetic diversity, MNTD.st is standardized mean nearest taxon distance, and MPD.st is standardized mean (phylogenetic) pairwise distance. We fitted generalised linear models (GLMs) and used model averaging to select the most parsimonious models. Only statistically significant predictors are shown. Points are mean coefficient estimates and filled curves are probability distributions of model coefficients.

**Figure 4.**
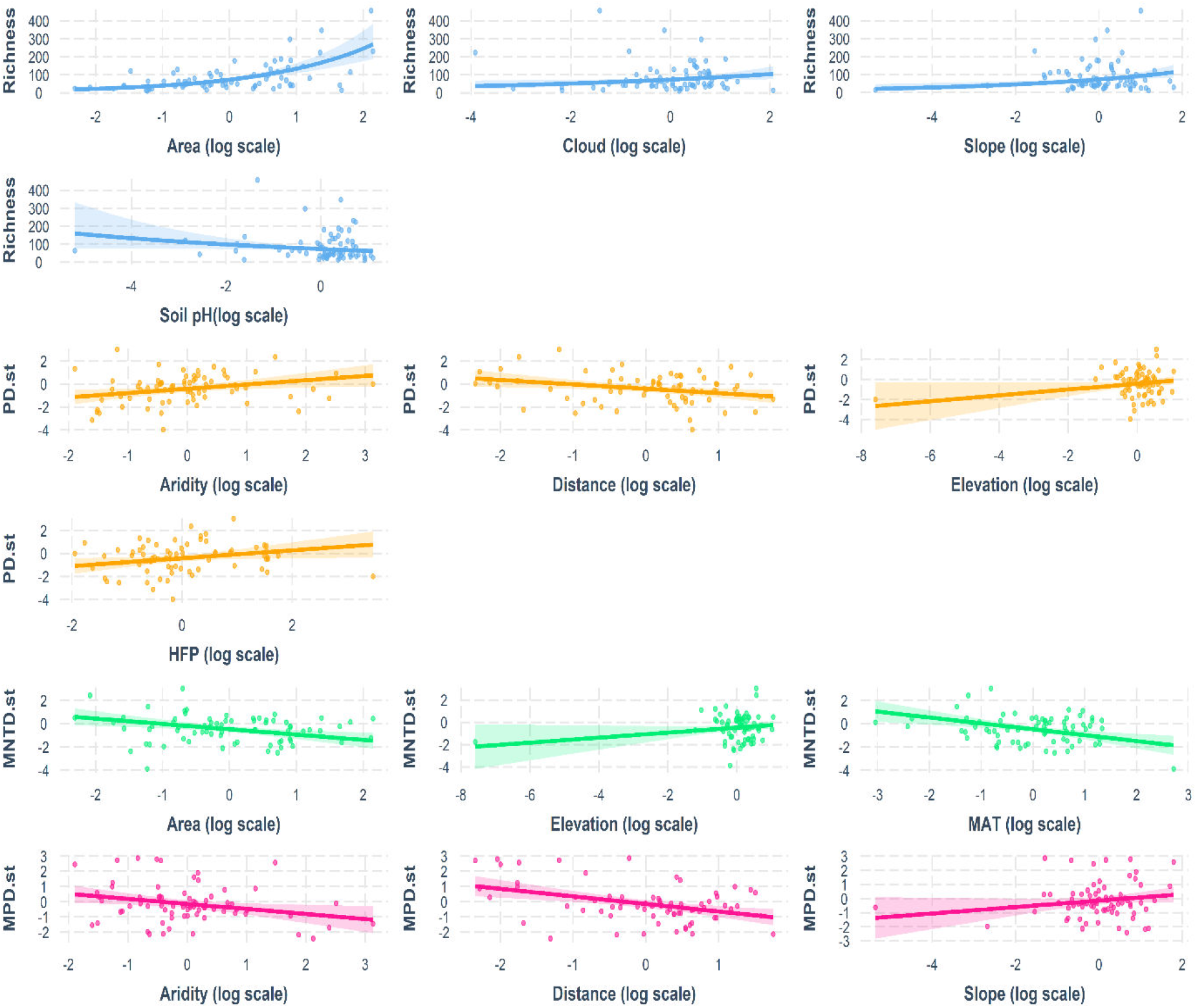
Estimated responses of taxonomic and phylogenetic diversity to environmental, topographic, and geographic factors across coastal lomas in Peru and Chile. HFP is the human footprint, MAT is mean annual temperature, and Distance is the distance of each coastal loma to the coast. We fitted generalised linear models (GLMs) and used model averaging to select the most parsimonious models. Solid lines are the fitted lines from generalised linear models and the shaded area represents the 95% confidence interval.

Finally, once accounting for the effects of climate, topography, and geography on taxonomic and phylogenetic diversity, we found no evidence of a latitudinal shift in taxonomic or phylogenetic diversity (p > 0.05; Figs. S3-6).

## 4. Discussion

Our results reveal that plant taxonomic and phylogenetic diversity of coastal lomas varies inversely and non-linearly with latitude, and is shaped by a combination of water availability, environmental heterogeneity, and distance from the coast. After accounting for these factors, we found no evidence that plant diversity of coastal lomas systemically shifts with latitude. The decline in species richness towards lower latitudes, has been reported in other taxa such as rocky intertidal (Thyrring & Peck, 2021), ants (Vasconcelos et al., 2017), mollusk species (Steffen & Sven, 2010), marine benthic algae (Santelices, 1980) and in ferns and mosses (Mateo et al., 2016).

Interestingly, this pattern has not been observed in vascular plants. The potential causes of inverse latitudinal gradients include more time for species to accumulate, faster diversification, more energy, and perhaps water availability (Hawkins et al., 2003), yet can be highly taxon-specific (Cerezer et al., 2022). Our finding supports the idea that latitude itself is not a direct, causal explanation for plant diversity on coastal lomas, but rather that the main environmental drivers (water availability) that vary with latitude, in addition to faster diversification rates (see Neves et al., 2021), are responsible for the latitudinal diversity gradient (LDG) in plants.

Futhermore, our results extend those of Manrique et al., (2014), who did not find a latitudinal gradient in plant species richness across a subset of 13 coastal lomas. Also, a previous study found that plant cover increases non-monotonically with latitude in *Tillandsia*-dominated coastal lomas (Pino et al., 2006), possibly in response to spatial variation in coastal fog and precipitation linked to ENSO events (e.g., Dillon, 2005).

While coastal fog productivity varies depending on proximity to the ocean, altitude, and local topography, it may not vary with latitude, at least in northern Chile (Larrain et al., 2022).

Our findings therefore highlight the importance of environmental variation at local spatial scales – and not macroecological scales- in determining diversity patterns in coastal lomas, which may have generated the non-linear latitudinal gradient that we observed for standardized Faith’s PD and MPD. Yet, we found that standardized MNTD exhibits a negative, linear latitudinal gradient, possibly suggesting that more recent diversifications have been constrained by water availability toward the northern extent of coastal lomas’ distribution. These can be explain by the onset of hyperarid conditions during the Pleistocene would have led to habitat fragmentacion, promoted plant speciacion (Merklinger et al., 2021). Examples of plant genera that have diversified during hyper-arid Peruvian Chilean coastal desert conditions include *Cristaria* (Malvaceae), which diversified around ∼3.7 ± 2 Ma (Boehnert et al., 2019), *Nolana* (Solanaceae) dating to ∼2 Ma (Guerrero et al., 2013), and *Eulychnia*, which diversified approximately 0.82 Ma (Merklinger et al., 2021). Also, low standardized MPD values at the extreme south and north of the coastal lomas’ distribution of appear to indicate stronger niche conservatism of coastal lomas plants at hight and low latitudes. The contrasting latitudinal patterns for phylogenetic diversity measures associated with different evolutionary time scales therefore suggest that species diversification events, together with historical changes in climatic conditions, may have shaped plant diversity patterns across coastal lomas in South America.

Overall, we found low values of unstandardized PD, which coincides with values reported for other drylands such as the Sonoran Desert, large parts of Caatinga and Atlantic forest (Amaral et al., 2022), and the Atacama Desert (Scherson et al., 2017). This suggests that strong lineage clustering, i.e., neoendemism, has shaped the composition of the flora of coastal lomas at macroecological scales. For example, *Tillandsia landbeckii,* a monospecific loma forming species in the Atacama Desert, is thought to have evolved only 2 mya (Möbus et al., 2021). Because of their relatively recent origins, most deserts may have relatively lower phylogenetic diversity when compared to relatively more humid environments (Scherson et al., 2014; Neves et al., 2021). Yet, overall phylogenetic endemicity in seed plants reaches its peak in the tropical regions where the Peruvian Chilean desert is located (Cai et al., 2023). Across coastal lomas, we found evidence of increasing phylogenetic clustering towards the North for standardized MNTD, yet only in the South for standardized MPD, which was phyllogenetically overdispersed at ∼22°S and 18°S. Together, the results of the phylogenetic metrics suggest that different processes may have operated at different evolutionary times along coastal lomas distribution. However, we found that the 95% confidence intervals of standardized PD overlapped with zero, suggesting that either neutral processes (Vellend, 2016) or weak niche partitioning and other biotic processes, such as competition (Cadotte & Davies, 2016) also may have shaped community assembly at local scales.

Our models showed that current environmental conditions, particularly those associated with water availability, have a significant effect on taxonomic and phylogenetic diversity across coastal lomas. Standardized PD Faith’s increased positively with aridity, which is broadly consistent with global and regional studies of drylands, desert scrub, and coastal lomas that reported increases in phylogenetic diversity with decreasing aridity (Berdugo et al., 2020; Stotz et al., 2021, Xiao et al., 2018). Moreover, the negative relationship between standardized MPD and aridity suggests that the diversity of older plant clades has been constrained by water availability (Neves et al., 2021). On the other hand, the positive relationship between aridity and standardized Faith’s PD may indicate high immigration or in-situ diversification rates in areas with high aridity (Fritz & Rahbek, 2021; Mazel et al., 2016). Yet, plant diversification in the Atacama Desert could have happened after the increase in aridity, which occurred in the late Pliocene after 5 Ma (Hartley & Chong, 2002). This diversification may be associated with pluvial events during the Pleistocene that may have facilitated the long-distance dispersal of Andean species to the coastal desert (Díaz et al., 2012). Conversely, the inferred onset and expansion of hyper-arid conditions towards higher elevations in the Precordillera and Pre-Andean basins (Hartley & Chong, 2002) likely restrict the distribution of plant populations in the Atacama Desert. Well-known examples include the genera *Nolana* and *Cristaria* (see Dillon et al., 2009; Boehnert et al., 2019). Further research on the dynamics of modern speciation events across coastal lomas is needed to strengthen our understanding of how temporal changes in aridity have impacted current diversity patterns.

Our results clarify the impacts of the dynamic interplay between environmental conditions and topography on diversity patterns in coastal lomas. Measures of environmental heterogeneity, such as slope and elevation, are frequently associated with diversity patterns (Heidrich et al., 2020; Stein et al., 2014), and also co-vary with climatic conditions, such as temperature and precipitation. In coastal lomas, slope and elevation are related to availability of coastal fog (see Muenchow et al., 2013; Rundel & Mahu, 1976) and have been found to influence taxonomic diversity at local scales (Moat el al., 2021). Furthermore, topographically heterogeneous areas should provide more possibilities for shelter and refuge, enabling species to persist in extreme conditions such as those found in deserts (e.g., Keppel et al., 2012).

We also found evidence that diversity patterns across coastal lomas depend on their biogeographical context. As expected, plant species richness increased positively with the area of coastal lomas, a trend previously documented for coastal lomas at more local scales (Arana, 2019). Moreover, standardized MNTD decreased with increasing area, suggesting a convergence in community composition in larger lomas. This convergence is characterized by species exhibiting similar life strategies and growth forms, which may have adaptations to local environmental conditions. Additionally, El Niño-Southern Oscillation (ENSO) events, which are known to enhance moisture levels and facilitate the expansion of plant distributions (Campbell, 1982), likely have affected the spatial extent of coastal lomas over time. Furthermore, decreases in coastal lomas area have been primarily associated with land-use change (González & Torres, 2009). However, human activity does not always have a predictable impact on plant diversity, as we found an increase in standardized PD with the human footprint index (HFP), possibly due to the introduction of exotic species that can positively affect diversity metrics.

We also observed that standardized MPD peaked at ∼22.5 and ∼13.5°S, which coincides with distinct geographical barriers, the Atacama Desert, Lima Chillon and Lurin hydrographic basins in the Center Peru and the Deflection of Abancay in Central Peru (Lima and Apurimac department). These geographical barriers could have influenced species dispersion and divergence, resulting in coastal lomas with a mixture of distantly related taxa (Mazel et al., 2016). Lastly, we found that standardized PD and MPD decreased with increasing distance from the coast, again highlighting the importance of coastal fog for maintaining species coexistence.

### Uncertainty of the present study

Although our dataset of coastal lomas is the most comprehensive compiled to date, it still may have spatial, phylogenetic, taxonomical and historical biases (Hortal et al., 2008; Hughes et al., 2021; Meyer et al., 2006) that may have effected our results.

Firstly, at least 30% of coastal lomas had no published species checklist, reflecting collectors bias. To complete our data set, we therefore used species records from other data sources, e.g., herbaria and GBIF, which may underestimate their true diversity (e.g., Hopkins, 2019). Secondly, most plant diversity data of coastal lomas is limited to conservation areas (Gonzales et al., 2023); by not including less protected coastal lomas, we may have underestimated human activity impacts on the taxonomic and phylogenetic diversity patterns of coastal lomas. Thirdly, the lack of high-resolution climate data on fog and low clouds along the South American coast limits our understanding of the role of both factors play in shaping the plant diversity of coastal lomas (see del Río et al., 2021). Lastly, we used a relatively coarse-scale resolution (1 km^2^) to estimate environmental conditions and environmental heterogeneity in coastal lomas. A higher resolution of Digital Elevation Model DEM data may better capture the environmental heterogeneity and micro-climate of coastal lomas, which may has been found to improve predictions of species distribution in other ecosystems (Lembrechts et al., 2019). Finally, global climatic data sets used in regional studies may fail to capture local climatic conditions, particularly in regions with a low density of climate stations and sharp climatic gradients as found in topographically complex areas (Bedia et al., 2013).

## 5. Conclusions

In this study, we collated the most comprehensive data set of coastal lomas to evaluate the underlying drivers of plant taxonomic and phylogenetic diversity in South America. We found that plant diversity (taxonomic and phylogenetic) exhibits an inverse latitudinal diversity gradient, which is largely attributable to a peculiar spatial variation in climate and topography. We provide evidence that drivers of biodiversity differ markedly among phylogenetic diversity indices, revealing how coastal lomas phylogenetic structure has shifted over evolutionary time. However, our understanding of the potential impacts of land-use change on plant diversity of coastal lomas remains limited, as there is little data on coastal lomas outside conservation areas in Perú and Chile. In view of our results, future research should focus on coastal lomas without checklist data (geographic shortfall; Hortal et al., 2008) to increase our ability to predict their response to global change drivers, particularly climate and land-use change. Furthermore, adaptation and planning strategies should be developed in response to the increasing aridification and land degradation in coastal lomas to conserve their unique biodiversity.

## Acknowledgements

We would like to thank to the New Flora of Chile project, the herbarium of the University of Concepción, and Diego Alarcón for extracting species records. We would also like to thank Dr. Asuncion Cano of the Universidad Mayor de San Marcos for providing access to the herbarium, Dr. Victor Quipuscoa for providing access to the facilities at the Michael Owen Dillon Institute IMOD. Dr. Patrick Weigelt, Dr. Amanda Taylor, Dr. Maria Fernanda Perez, Dr. Christian Hernandez, Dr. Claudio Latorre, Dr. Miguel Fariña, and Maria Laura Tolmos provided valuable suggestions that substantially improved the manuscript. Finally, we would thank to Dr. Juan Armesto for his advice and guidance during the PhD of FG. This work was supported by the E032-117-2017-Doctorate scholarship grants PROCIENCIA/CONCYTEC-PERU.

## 6. Appendices

Supplementary Information.

## Appendices

Supplementary Material

## **Appendix S2**. Macroecological hypotheses evaluated to explain plant diversity patterns across drylands, associated environmental predictors and references

| Hypotheses | Prediction | Variable | Reference |
| --- | --- | --- | --- |
| Species–energy relationship (SER) | Positive | temperature media anual (Bio1), net primary productivity (NPP), aridity, isothermality (Bio3) | Currie (1991) |
| Water–energy dynamics (WED) | Positive | annual cloud cover, cloud cover for August, precipitation of the wettest month (Bio13), annual precipitation (Bio12), precipitation of the driest month (Bio14), mean monthly climate moisture index (cmi_mean), | Hawkins et al. (2003) |
| Environmental heterogeneity | Positive | Aspect (sino and cosine), topography, roughness, elevation, slope | Pianka (1966), Stein et al., (2014) |
| Species-area | Positive | Area (km2) | Rosenzweig (1995) |

**Figure S1.**
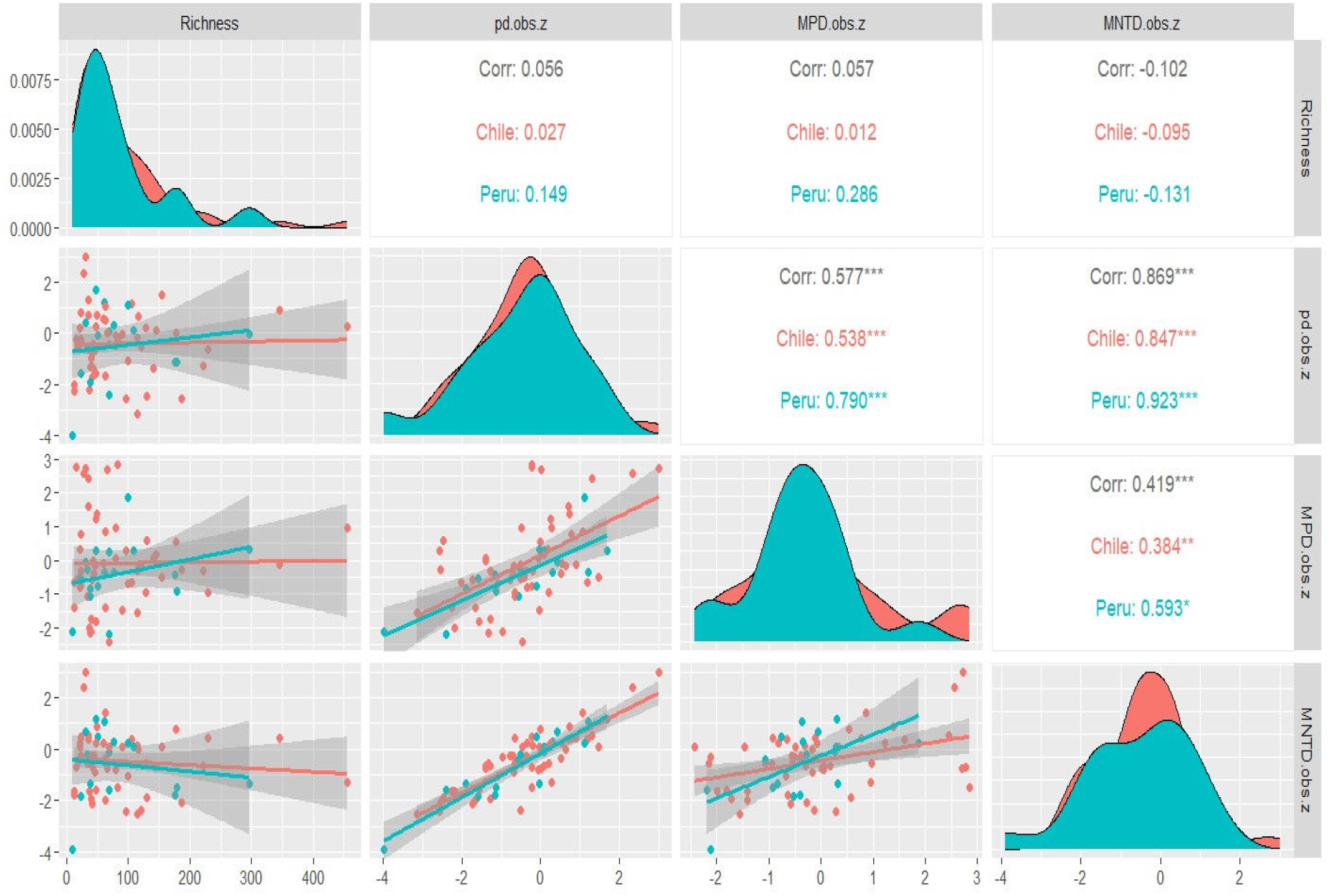
Pearson correlations between richness and standardized phylogenetic diversity indices (PD.st: pd.obs.z, MPD.st: MPD.obs.z, MTND.st; MNTD.obs.z). Red corresponds to coastal lomas in Chile, while blue corresponds to those in Perú.

**Figure S3.**
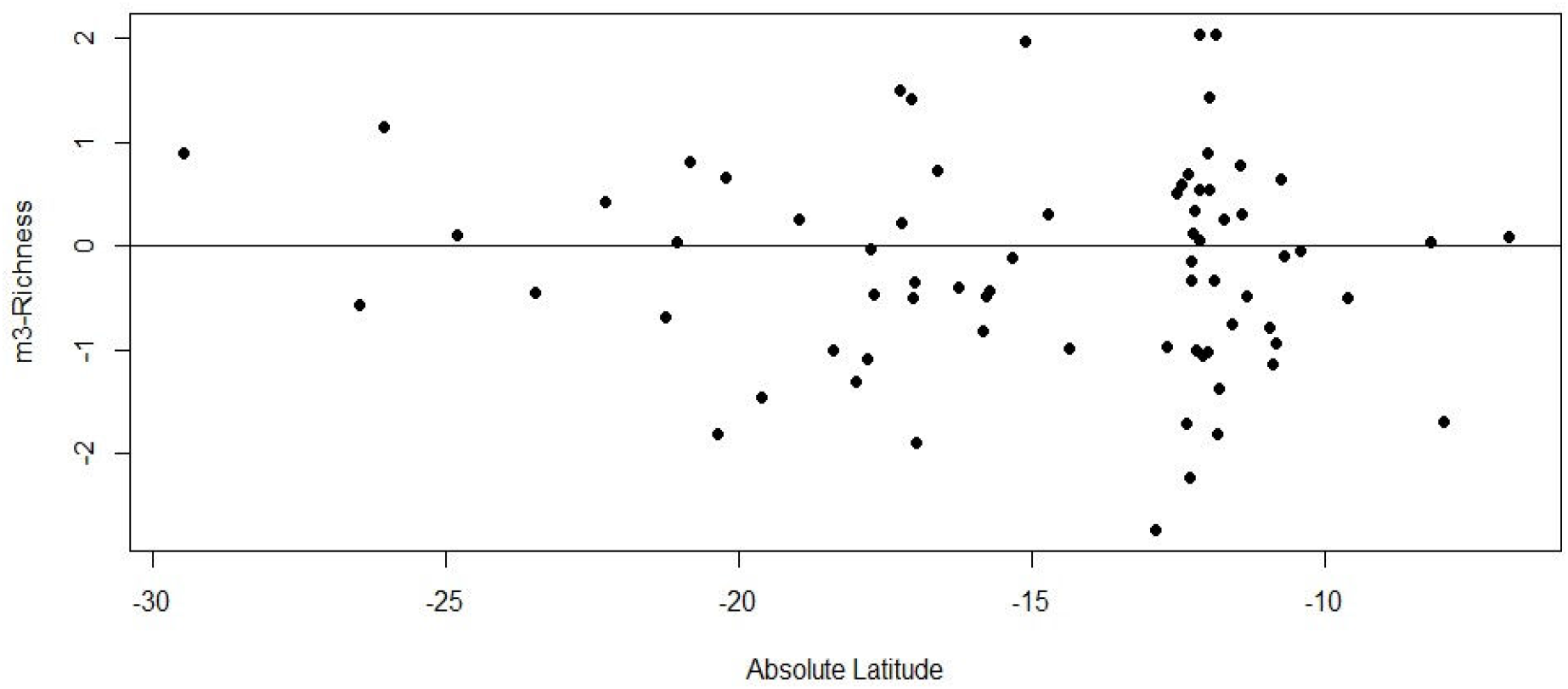
Relationship between residuals of the species richness model and absolute latitude.

**Figure S4.**
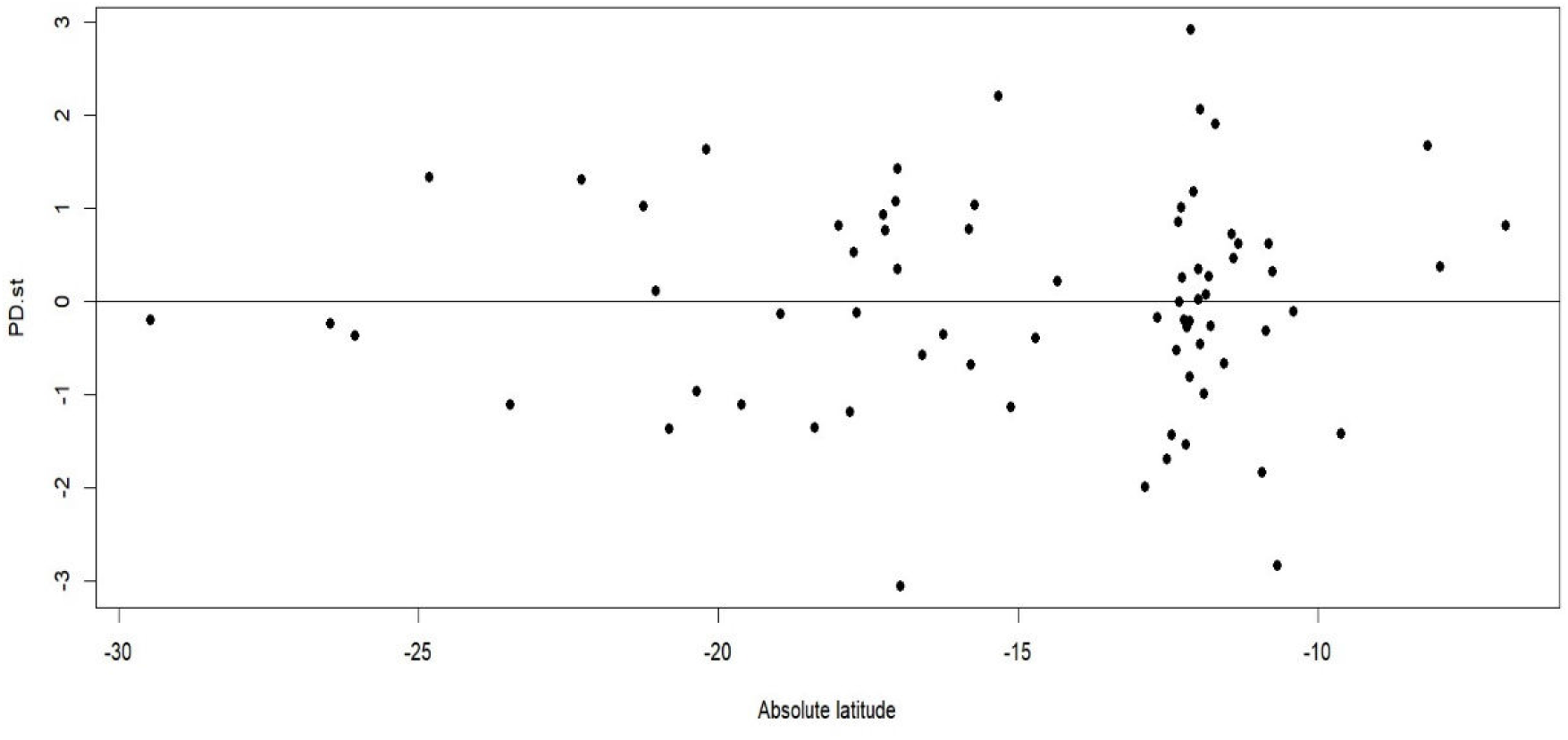
Relationship between residuals of the standardized phylogenetic diversity (PD.st) model and absolute latitude.

**Figure S5.**
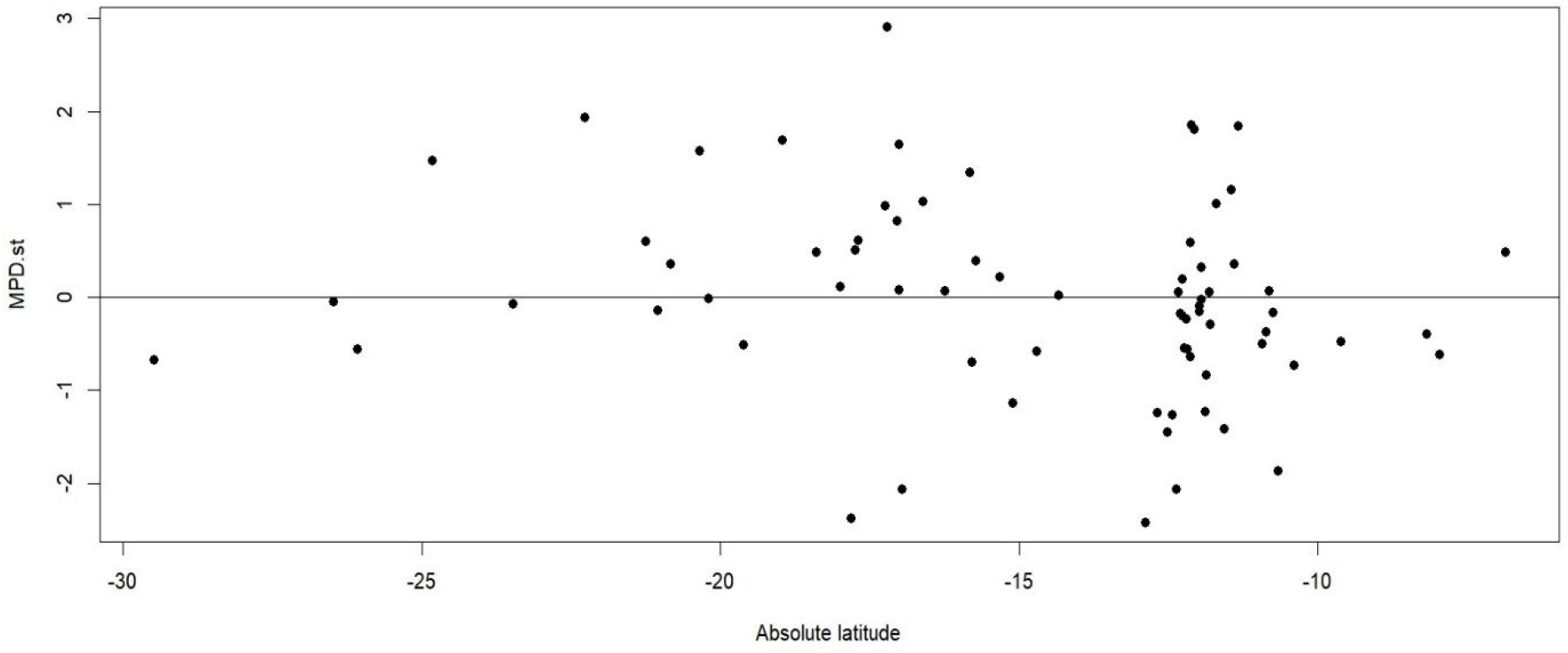
Relationship between residuals of the standardized Mean Pairwise Distance (MPD.st) model and absolute latitude.

**Figure S6.**
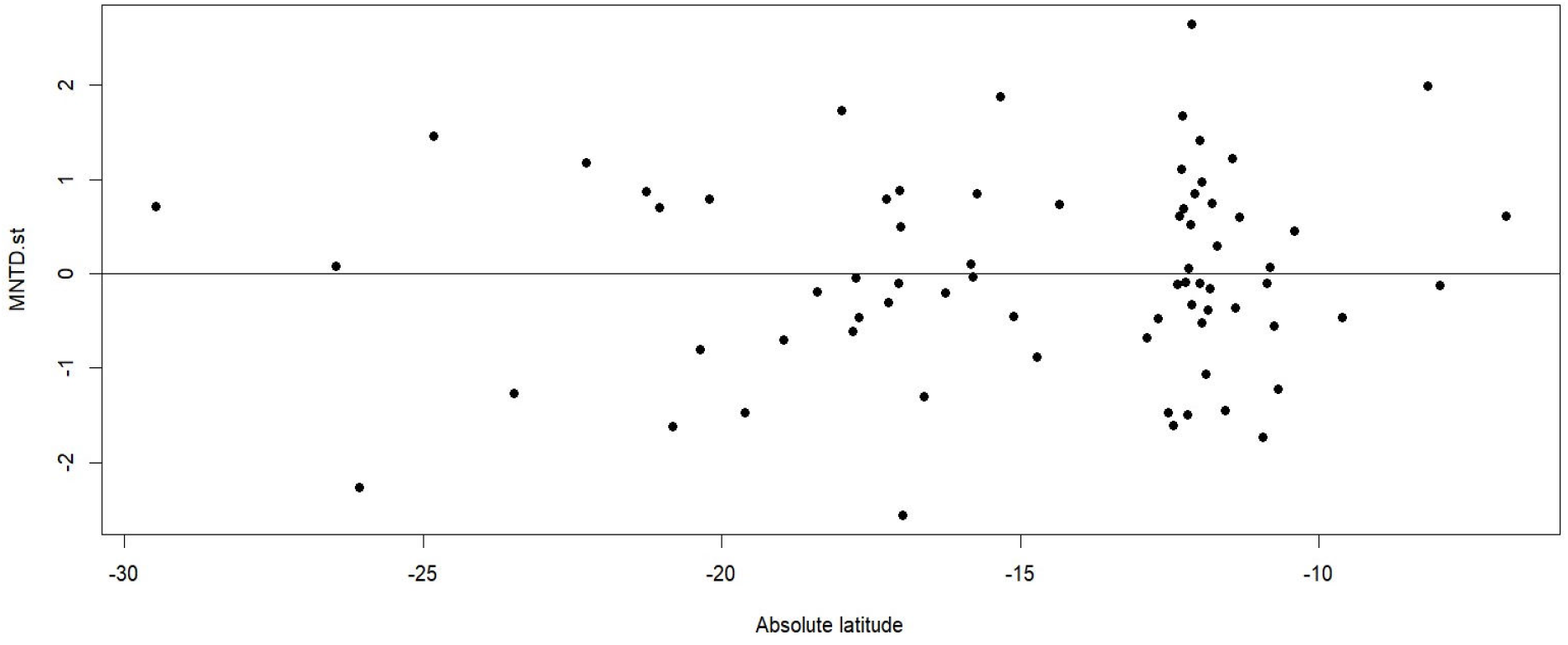
Relationship between residuals of the standardized Mean Nearest Taxon Distance (MNTD.st) model and absolute latitude.

**Table S1.** Generalised lineal models (GLM) evaluating the response of plant species richness and standardized phylogenetic diversity to environmental, geographical and topographical variables in coastal lomas across Perú and Chile. We performed model selection using AIC to identify the most parsimonious model for each response variable.

|  | Species richness | PD.st | MNTD.st | MPD.st |
| --- | --- | --- | --- | --- |
| Area | 0.61***<br>(0.08) |  | -0.46**<br>(-0.15) |  |
| Cloud_m | 0.18<br>(0.14) |  |  |  |
| Slope | 0.24**<br>(0.07) |  |  | 0.23<br>(0.12) |
| Soil (pH) | -0.15<br>(0.08) |  |  |  |
| Aridity |  | 0.37*<br>(0.18) |  | -0.33*<br>(0.16) |
| Elevation |  | 0.30<br>(0.25) | 0.22<br>(0.15) |  |
| Distance |  | -0.38*<br>(0.15) |  | -0.50**<br>(0.16) |
| HFP |  | 0.34*<br>(0.15) |  |  |
| MAT |  |  | -0.51**<br>(0.18) |  |
| N | 72 | 72 | 72 | 72 |
| AIC | 738.32 | 234.30 | 219.53 | 222.72 |
| BIC | 751.98 | 247.96 | 230.91 | 234.10 |
| Pseudo R<sup>2</sup> | 0.062 | 0.2 | 0.21 | 0.28 |

Standard errors are robust to heteroscedasticity. \*\*\* $p < 0.001$ ; \*\* $p < 0.01$ ; \* $p < 0.05$ .

